# *Pseudohydrosme bogneri* sp. nov. (Araceae), a spectacular Critically Endangered (Possibly Extinct) species from Gabon, long confused with *Anchomanes nigritianus*

**DOI:** 10.1101/2021.03.25.437040

**Authors:** Lois Moxon-Holt, Martin Cheek

## Abstract

A Gabonese taxon cultivated for decades in the botanic gardens of Europe as *Anchomanes nigritianus* is shown to be a new species to science, and on current evidence, is best placed as the fourth species of the Gabonese-centred, poorly known genus *Pseudohydrosme*. Data on the morphological separation between *Anchomanes* and *Pseudohydrosme* are reviewed. Although phylogenomic studies may show in future that the two genera need to be merged, for the moment their separation is reinforced on morphological grounds. *Anchomanes* lacks the spathe tube, ovoid-globose, 2 – 4 locular pistil and thick, lobed stigma on a symmetric, stout style that we show to characterise the redelimited *Pseudohydrosme*. (*Anchomanes* has oblong, polygonal, 1-locular pistils, stigmas asymmetric, sessile, thin and disc-like or on asymmetrical conical styles and are pointed or brush-like). In addition, *Pseudohydrosme* (where known) has stipitate (versus sessile) fruits and on current evidence lacks the lacticifers recorded from *Anchomanes*. We test the hypothesis that the taxon is a new species to science, naming it as *Pseudohydrosme bogneri*, and conclude that it is Critically Endangered (Possibly Extinct) using the IUCN 2012 standard. *Pseudohydrosme bogneri* appears to be the tenth documented probable global extinction of a plant species that has occurred among the narrowly endemic plant species of the Libreville area, Gabon.

## INTRODUCTION

In 1973, the late great aroid specialist Josef Bogner (Renner & Mayo 2020) visited Gabon to look for the long-lost genus *Pseudohydrosme* Engl. (Engler 1892) (Bogner 1981). He succeeded in finding one of the two species then known of the genus, *Pseudohydrosme gabunensis* Engl., and brought live material back to Munich Botanical Garden for cultivation, in addition to preserved material. Bogner also collected other aroids and plants of other families several of which were later published as new species to science e.g. *Palisota bogneri* Brenan (Brenan 1984) and *Rhaphidophora bogneri* P.C. Boyce & Haigh (Boyce & Haigh 2016).

Bogner found *Pseudohydrosme gabunensis* in a remnant (c. 400 m x 400 m) of original forest that survives in the capital city Libreville, now known as the Sibang arboretum. In his ecological notes to his paper on that species, he states “other plants found together are *Anchomanes nigritianus* Rendle, a very rare species first found in South Nigeria, of which I collected one specimen only at a distance of less than 100 m from the locality of *P. gabunensis*. Also very common were *Anchomanes difformis* (Blume) Engl., *Amorphophallus maculatus* N.E. Br., which grew nearby. *Pseudohydrosme gabunensis* flowers before the single leaf appears and I found it flowering in October, whereas *Anchomanes* normally flowers together with the leaf….”. He illustrates as Figure 13 *Anchomanes nigritianus* with a photo of the inflorescence and the petiole giving the collection as *Bogner* 662 (Bogner 1981:37). This plant was also taken back to Munich Botanical Garden and successfully cultivated, apparently producing inflorescences and leaves in subsequent years (Fig. 1 & 2). Six herbarium specimens, three with leaves, three with inflorescences are preserved at M. Different sheets have labels in different formats and descriptions, and are written in different hands, giving support to the idea that they were made at different times although all were given the specimen number *Bogner* 662. Additional inflorescences and leaves were sent as specimens to US and K, where they are also preserved under this number.

**Fig. 1.**
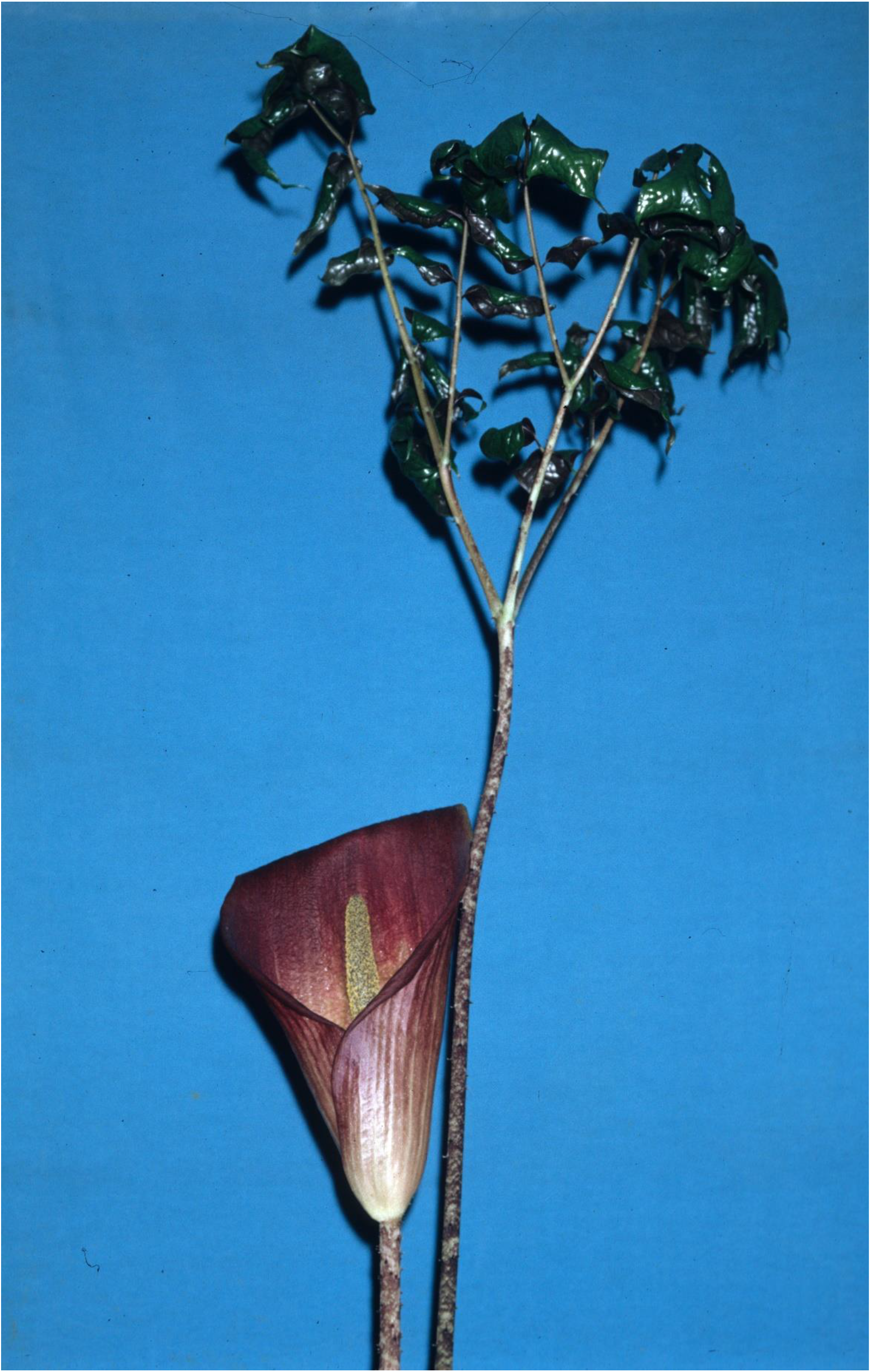
Pseudohydrosme bogneri. Habit, plant with flower at anthesis and leaf-blade unfurling. Note the long petiolules of the proximal leaflets. Cultivated at Munich Botanic Garden, *Bogner* 662. Photo Jan. 1995 by Günter Gerlach

**Fig. 2.**
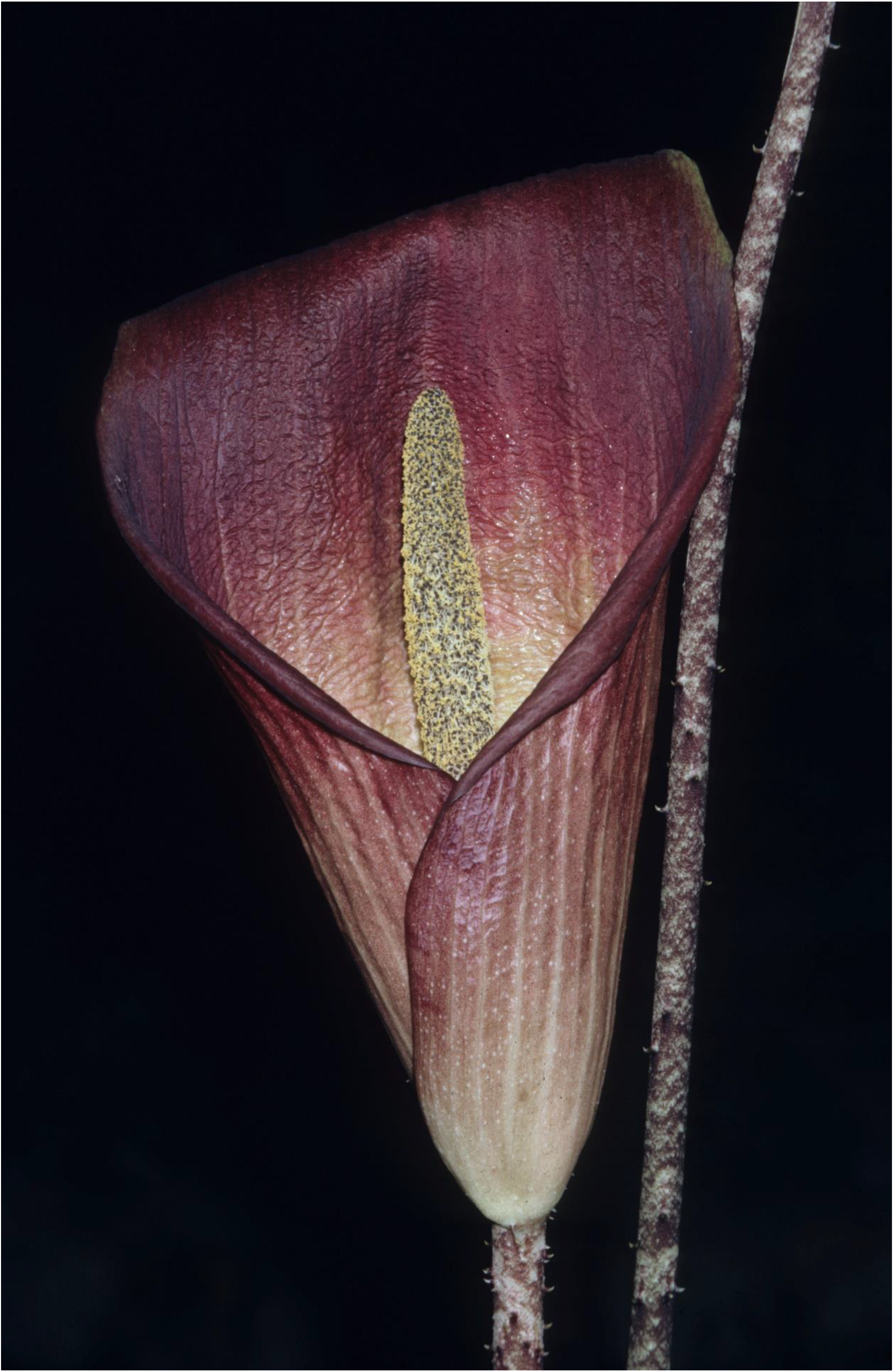
Pseudohydrosme bogneri. Close-up of inflorescence in late anthesis (male phase). Cultivated at Munich Botanic Garden, *Bogner* 662. Photo Jan. 1995 by Günter Gerlach

Bogner had collected a second plant of this species, also from Sibang, collected as a juvenile. This was numbered *Bogner* 640 (K000499336 http://www.plantsoftheworldonline.org/taxon/urn:lsid:ipni.org:names:84501-1). He also brought this back to Europe as live material and it may have been when it flowered that he identified it as the same taxon as *Bogner* 662. Flowering material of *Bogner* 640 is preserved in spirit at K in addition to *Bogner* 662. Bogner initially identified these plants as *Anchomanes giganteus* Engl. (a species described from the Democratic Republic of Congo, DRC) but three years later, in 1976, he re-annotated the herbarium specimens (see link above) as *A. nigritianus* Rendle, a species previously known only from the type specimen collected in Nigeria.

Bogner’s identification has not been tested until now. His material of *Bogner* 662 and *Bogner* 640, is figured as *Anchomanes nigritianus* in Mayo *et al.* (1997), and plants thought to be from this source, cultivated by Wilbert Hetterscheid in the Netherlands, which are consistent with the preserved material, are also available as images on the website of the International Aroid Society (http://www.aroid.org/genera/speciespage.php?genus=anchomanes&species=nigritianus Accessed 21 March 2021). However, in the course of a research project revising the genus *Anchomanes*, a comparison of the dimensions and proportions of the inflorescence of Bogner’s Gabonese material with that of the only material known of *Anchomanes nigritianus*, the type from Oban, Nigeria (*Talbot* 1247, BM), shows morphological differences so numerous and disjunct that they cannot be accommodated in the same species (see Table 1 below).

**Table 1.** Diagnostic characters separating *Anchomanes nigritianus* Rendle from *Pseudohydrosme bogneri* sp. nov. Data for the first species from Rendle (1913) and from the type specimen *Talbot* 1247 (BM, photo M!). Data for the second species is taken from *Bogner* 662 (K, M, US) and *Bogner* 640 (K).

|  | *Anchomanes nigritianus* | *Pseudohydrosme bogneri* |
| --- | --- | --- |
| Length of spathe | 40 cm | 27 – 30 cm |
| Length of spadix | 8 – 10 cm | 18 – 22.5 cm |
| Ratio of lengths<br>spadix:spathe | 1:4 | 1:1.4 – 1.8 |
| Shape of ovary | obovoid | ovoid-globose |
| Length of pistil | 2 – 2.5 mm | 5.5 – 6.5 mm |
| Width of stigma | 2 mm | 3 – 4.5 mm |
| Stigma lobing | unlobed | 2 – 3-lobed |
| Pistil:number of locules and<br>ovules | 1 | 2 – 3( – 4) |
| Distribution | Oban, S.E. Nigeria | Libreville, Gabon |
| Spadix, length of ♀ zone | 4 – 5 cm | 8 – 11 cm |
| Spadix, length & colour of<br>♂ zone | 4 – 5 cm ‘white-pale bluish<br>purple (‘mauve’ or ‘lilac’) | 8 – 11 cm blackish purple |

Moreover, re-examination of Bogner’s material shows it to have the key features of *Pseudohydrosme*, a genus of three species closely related to *Anchomanes* but centred in Gabon, and which was recently revised (Cheek *et al.* 2021). In this paper we describe morphologically the taxon represented by Bogner’s material for the first time, and we test the hypothesis that it is a fourth, hitherto undescribed species of *Pseudohydrosme*. We propose the name *Pseudohydrosme bogneri* Cheek & Moxon-Holt sp. nov. for this taxon.

## METHODS

Herbarium citations follow Index Herbariorum (Thiers *et al.* 2020). Specimens were viewed at B, BR, K, M, MA, MO, P, US, WAG. The material that is the subject of this paper, and the genus *Pseudohydrosme* are centred in Gabon. The national herbarium of Gabon is LBV, but the most comprehensive herbaria for herbarium specimens of that country are P and WAG. During the time that this paper was researched, it was not possible to obtain physical access to material at WAG (due to the transfer of WAG to Naturalis, Leiden, subsequent construction work, and covid-19 travel and access restrictions). However, images for WAG specimens were studied at https://bioportal.naturalis.nl/?language=en and those from P at https://science.mnhn.fr/institution/mnhn/collection/p/item/search/form?lang=en_US. Additionally, we received digital images that we requested from M (Bogner’s herbarium) and, regarding Equatorial Guinea, neighbouring Gabon, from MA. We also searched JStor Global Plants (2020) for additional type material of the genus, and finally the Global Biodiversity Facility (GBIF, www.gbif.org accessed March 2021) which lists 28 occurrences and 21 images of *Pseudohydrosme*, mainly relating to the holdings (including duplicate herbarium sheets) of WAG, followed by P.

Binomial authorities follow the International Plant Names Index (IPNI 2020). The conservation assessment was made using the categories and criteria of IUCN (2012). Spirit preserved material was available at K. Herbarium material was examined with a Leica Wild M8 dissecting binocular microscope fitted with an eyepiece graticule measuring in units of 0.025 mm at maximum magnification. The drawings were made with the same equipment using Leica 308700 camera lucida attachment. This was used to characterise and measure particular features of the flowers. The new species described below as *Pseudohydrosme bogneri* was mainly described from herbarium specimens (gross morphological structures) and spirit preserved material (details of the spadix). The terms and format of the description follow the conventions of Mayo *et al.* (1997) and Cheek *et al.* (2021). Georeferences for specimens lacking latitude and longitude were obtained using Google Earth (https://www.google.com/intl/en_uk/earth/versions/). The map was made using SimpleMappr (https://www.simplemappr.net).

## RESULTS

The diagnostic differences between *Anchomanes nigritianus* and *Pseudohydrosme bogneri* are given in table 1 below. The first species remains very imperfectly known.

Transverse sections of spirit-preserved material of female flowers of the new species showed that the number of locules per pistil varied from 2 – 4. In *Bogner* 640 (K spirit) two flowers were 2-locular, two were 3-locular and one was 4-locular. In *Bogner* 662 (K spirit) five were 2-locular and one was 3-locular.

### Generic placement: *Pseudohydrosme* versus *Anchomanes*

According to Mayo *et al.* (1997:221) and Hetterscheid & Bogner (2013), the only clear-cut distinction between *Anchomanes* and *Pseudohydrosme* is locule number. *Anchomanes* are 1-locular, and *Pseudohydrosme* 2(– 3)-locular. However, in a recent taxonomic revision of *Pseudohydrosme*, five additional characters separating the genera are given (Cheek *et al.* 2021, Table 2, see discussion below). In one of these, peduncle length, the new species described here fits *Anchomanes*, not *Pseudohydrosme*. In a second of these five additional characters (spadix: spathe proportions and the presence of a long spathe tube), the new species seems closer to *Anchomanes* than to *Pseudohydrosme* in spadix length, but not in spathe tube features. In some respects, therefore, this taxon seems intermediate between the two genera morphologically, supporting the suggestion that they be merged. On the other hand, several of the traits currently attributed to *Anchomanes* (e.g. in Mayo *et al.* 1997) were due to the inclusion of *Bogner* 640 & 662. If these are subtracted from the current delimitation of that genus, the morphological separation between the two genera needs to be reinterpreted.

**Table 2.** Characters separating *Anchomanes* and *Pseudohydrosme*. Revised and updated from Cheek *et al.* (2021).

|  | *Anchomanes* | *Pseudohydrosme* |
| --- | --- | --- |
| Spathe tube | Absent | Present, forming the proximal 1/3-2/3 of the spathe |
| Pistil shape | Oblong, polygonal | Globose or broadly ovoid, non-polygonal |
| Ovary locularity and stigma lobe number | 1 | 2 -3(-4) |
| Stigma & Style | Stigma sessile, thin, brush-like or pointed, never lobed (but can split on drying in *Anchomanes welwitschii*)<br>Style absent or asymmetrically conical | Stigma thick, lobed, lobes reflexed<br>Style cylindric, stout, symmetrical |
| Berries | Sessile | Stipitate (where known) |
| Laticifers | Present, simple, articulated | Absent (not detected by Keating 2002) |

When this is done, we can see that the disjunctions in morphology of the female flower between the redrawn genera are much larger than previously realised. Shorn of *Bogner* 640 and 662 (“*Anchomanes nigritianus*”), *Anchomanes* species now all possess, without exception (on current evidence) polygonal, oblong-obovoid pistils which either lack completely both a style and thickened stigma (the East African species *A. abbreviatus* Engl. and *A. boehmii* Engl. where the stigma is sessile, thin, disc-like, completely flat and asymmetrically placed (Mayo *et al.* 1997: Plate 70V&Z)) or where the style-like top of the pistil is conical, asymmetric and terminates in either a minute inconspicuous stigma, or a brush-like one (the Guineo-Congolian species, e,g, *A. difformis, A. welwitschii* Rendle (Mayo *et al.* 1997: Plate 70K, CC, FF)). In contrast in all four species of *Pseudohydrosme*, the ovary is globose or globose-ovoid, and not polygonal. The style is always symmetrical, distinct, stout, cylindric and bears a thickened, 2 – 3-lobed stigma, and the stigmatic lobes are reflexed (Mayo *et al.* 1997: Plate 70R, Plate 71H&I, Cheek *et al.* 2021 Fig.6G, Fig. 7E). Below, in Table 2, we redraw the morphological characters separating the two reinterpreted genera in the light of placement of *Bogner 640 and 662* in *Pseudohydrosme* rather than *Anchomanes*.

With this redelimitation, both genera are morphologically more coherent and more amply separated from each other than previously. The question will be further resolved by an imminent planned molecular phylogenomic study. For the moment, we describe the new species in *Pseudohydrosme* but accept that, depending on the outcome of the phylogenomic study, it may need to be transferred to *Anchomanes* in the future.

### Transition from *Pseudohydrosme* to *Anchomanes*

Images of stages of flowering of *Pseudohydrosme bogneri* taken by Wilbert Hetterscheid show that at early or pre-anthesis, the inflorescence (excepting the long peduncle) resembles that of *Pseudohydrosme:* the spathe is strongly fornicate, the distal part arching over the rest of the spathe, while the spathe tube, more than half the length of the spathe, completely conceals the spadix (http://www.aroid.org/genera/serveimage.php?key=892, accessed 21 March 2021). However, as anthesis proceeds, by the time that the male phase is reached, and pollen is produced, the apex of the spathe becomes reflexed leaving the tube uncovered, the spathe margins become revolute, the spathe opens out, unfurling at the base, reducing the length of the tube and exposing the spadix, in which phase the inflorescence more resembles *Anchomanes* (http://www.aroid.org/genera/serveimage.php?key=891 accessed 21 March 2021).

### Identification key to the sections and species of *Pseudohydrosme* (updated from Cheek *et al.* 2021)

1.Spadix with distal half covered in sterile male flowers. **Sect. *Zyganthera……P. buettneri***

1.Spadix lacking sterile flowers, distal part with fertile male flowers only. …………**Sect. *Pseudohydrosme*………2**

2.Peduncle over 1 m long, conspicuous; leaf unfurling at same time as flower produced; spadix about 2/3 the length of the spathe, the tip of the spadix protruding from the spathe tube in late anthesis…………***P. bogneri* sp. nov**

2. Peduncle < 0.05 m long, concealed by cataphylls; leaf not produced until at least weeks post-flowering; spadix about 1/4 the length of the spathe, completely concealed within the spathe tube…………3

3. Male and female zones of spadix contiguous; entire spadix covered in flowers densely arranged in both male and female zones; spathe blade inner surface yellow, greenish yellow or white with abrupt transition to a central dark red area; stigmas 2(–3)-lobed. Gabon (probably Congo-Brazzaville)…………***P. gabunensis***

3. Male and female areas of spadix incompletely contiguous; female flowers laxly arranged with axis of female zone partly naked especially distally; spathe blade inner surface light reddish brown or pink, with wide green veins, very gradually becoming darker towards the centre; stigmas 3(–2)-lobed. Cameroon…………***P. ebo***

### Pseudohydrosme bogneri *Cheek & Moxon-Holt* sp. nov. Type

Gabon, Libreville, Sibang Arboretum, fl. Oct. 1973, cultivated in Munich Botanical Garden, *Bogner* 662 (Holotype: K03822705; isotypes: K03822706, spirit K37350.000, M six sheets, US03822705, US03822706)

Diagnosis: differing from all other species of *Pseudohydrosme* Engl. in the peduncle exceeding 1 m in length, far exceeding the length of the spathe (versus peduncle only c. 1/10 the length of the spathe) and in the spadix protruding from the spathe tube in late anthesis (versus concealed within the spathe tube).

#### Perennial rhizomatous herb

Rhizome data not available. Leaf 1.25 – 1.35 m tall, petiole terete, 1.5 cm diameter at base, with large patent prickles up to 4 mm long. Blade of juvenile sagittate-elliptic 21.5 x 12 cm, adaxial surface silvery gray, abaxial surface deep wine red (label data from *Bogner* 640, K, http://specimens.kew.org/herbarium/K000499336) apex acute, basal lobes acuminate, basal sinus 9 x 11 cm, petiole 15 cm long. *Blade* of mature leaves dracontoid, divided into three main segments, each further divided once dichotomously, primary division 30 – 42 cm long, secondary divisions 22.5 – 34 cm long, lobes 11 – 16, usually pinnate sometimes bipinnate, dimorphic; larger, mainly distal lobes trapezoid 11 – 16 x 7-17 cm, apex truncate-bifid (biacuminate), acumen 1.2-2 cm long, decurrent to subsessile; smaller, mainly proximal lobes ovate, 7 – 11 x 3.5 – 6.5 cm, apex acuminate, base sometimes asymmetric, petiolules 0.5–2.5 cm long; lateral veins 3 – 5 on each side of the midrib, mostly running into margin, sometimes forming irregular submarginal vein, higher order veins reticulate. *Inflorescence:* Peduncle 105 – 120 x 1.5 cm, base purple-brown with olivaceous tinge, apex fawn brown (label data from *Bogner* 640, K) with large patent prickles up to 3(– 4) mm long, each prickle with dark purple mark below and white patch above (label data from *Bogner* 640, K). Spathe large, unconstricted, infundibuliform, 27 – 30 x 10 – 12 cm, 20 cm broad at widest point when opened out (label data from *Bogner* 662, M), basal half convolute, forming a funnel-shaped tube, limb comprising the distal half of the spathe, at anthesis inflexed over spadix (fornicate), later becoming reflexed, blade margins becoming revolute, apex acuminate; outer and inner surface purple-maroon but yellowing towards the base of the interior, with yellow longitudinal veins (label data from *Bogner* 662, M and *Bogner* 640, K), inner surface puckered (label data from *Bogner* 640, K). Spadix subcylindrical, 18 – 22.5 cm long. Female zone 8 – 11 cm long, 1.2 – 2 cm wide, contiguous with the male zone, moderately densely arranged distally at the apex, laxly arranged proximally, the spadix axis clearly visible between the flowers; stipe (spadix axis naked at the base), 1.5 – 2 cm long. Male zone 8 – 11 cm long, 1.7 – 3 cm wide narrowest at junction with the female zone, widest in the distal portion (Fig. 3B), apex rounded, completely covered in densely arranged male flowers, individual flowers impossible to distinguish, sterile flowers or appendix absent.

**Fig. 3.**
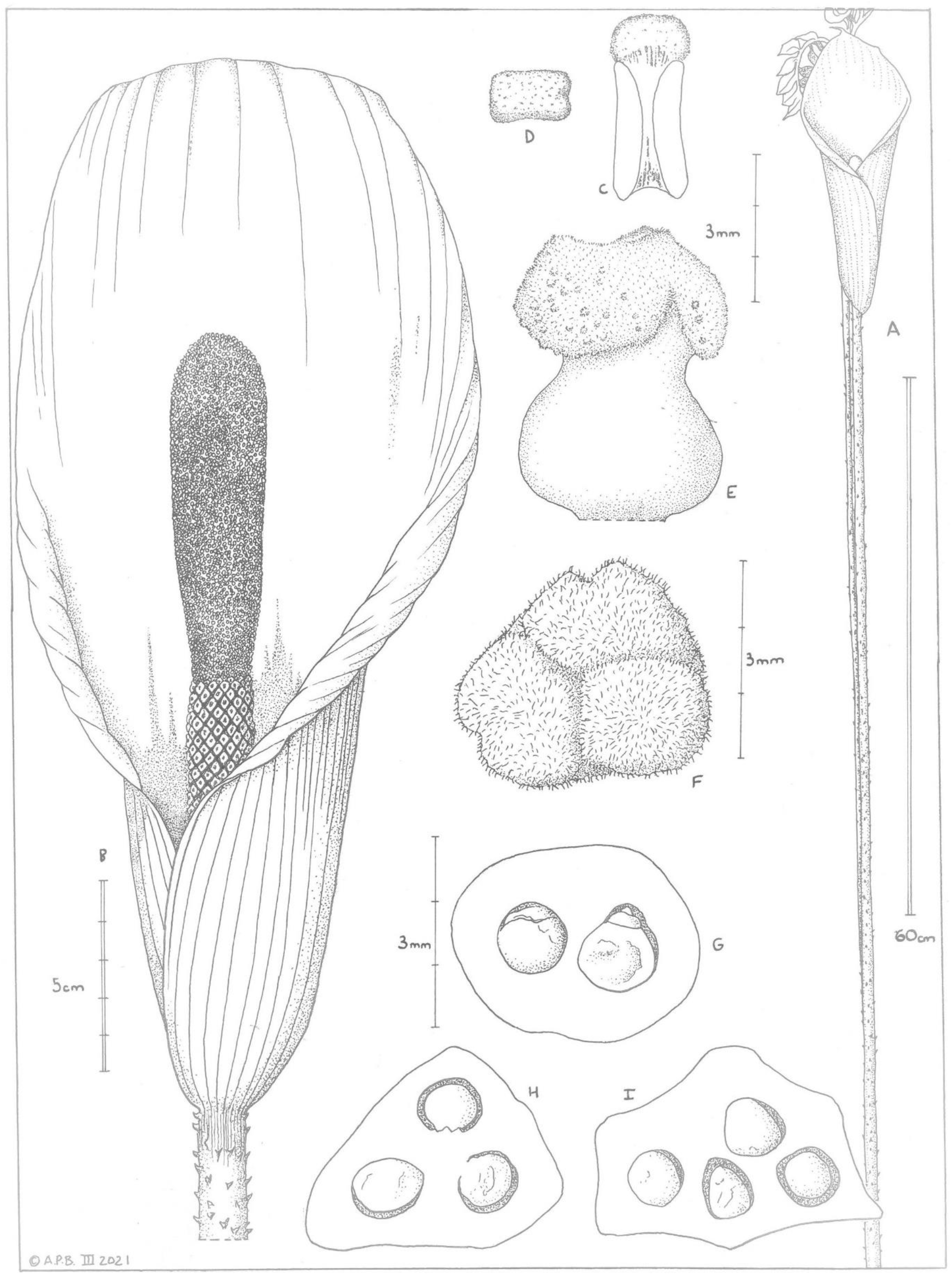
Pseudohydrosme bogneri. A habit, peduncle and inflorescence (unfurling leaf behind); B inflorescence showing spathe and spadix (spathe opened slightly to show female zone); C stamen, longitudinal section; D stamen, plan view; E pistil (female flower) side view; F trilobed stigma from above; G 2-locular ovary, transverse section; H 3-locular ovary, transverse section; I 4-locular ovary, transverse section. A from photo by Wilfred Hetterscheid (International Aroid Society); B-E from *Bogner* 640, (Kew spirit collection 29047.1) and *Bogner* 662 (Kew spirit collection 37350) after E. Catherine in Mayo, Bogner and Boyce (1997); F, H-I from *Bogner* 640 (Kew spirit collection 29047.1); G from *Bogner* 662 (Kew spirit collection 29047.1). Drawn by ANDREW BROWN.

#### Male flowers

with stamens sessile, prismatic, 3 – 4.3 mm long, in plan-view oblong 1.3 x 1.7 mm (Fig. 3D), apex convex, anther thecae lateral oblong, 2.7 – 3 mm long, overtopped by thick dark purple connective (label data from *Bogner* 640, K), pollen golden yellow, in strings (label data from *Bogner* 662, M) (Fig. 3C).

#### Female flowers

with ovary ovoid-globose, smooth, light green (label data from *Bogner* 662, M) 2.1 – 4.3 mm long, 3.1 – 4.3 mm diam., 2 – 3 – (4)-locular, style purple (label data from *Bogner* 662, M), 0.2 – 0.8 mm long, 2 – 2.5 mm diam., stigma brown purple (label data from *Bogner* 662, M), surface papillose and minutely hairy, 3–4.5 mm wide, 2 – 3 – (4)-lobed (Fig.3G-I), 0.75 – 1 mm thick; 2-lobed stigma with a small central depression and two raised peaks where the lobe margins meet; 3-lobed stigma with a central groove in each lobe, lobe apices rounded and reflexed. *Berry and seed* not seen.

## DISTRIBUTION

Gabon (Map 1). Restricted on current evidence to Libreville, Sibang Arboretum (possibly extinct, but see notes below).

**Map 1.**
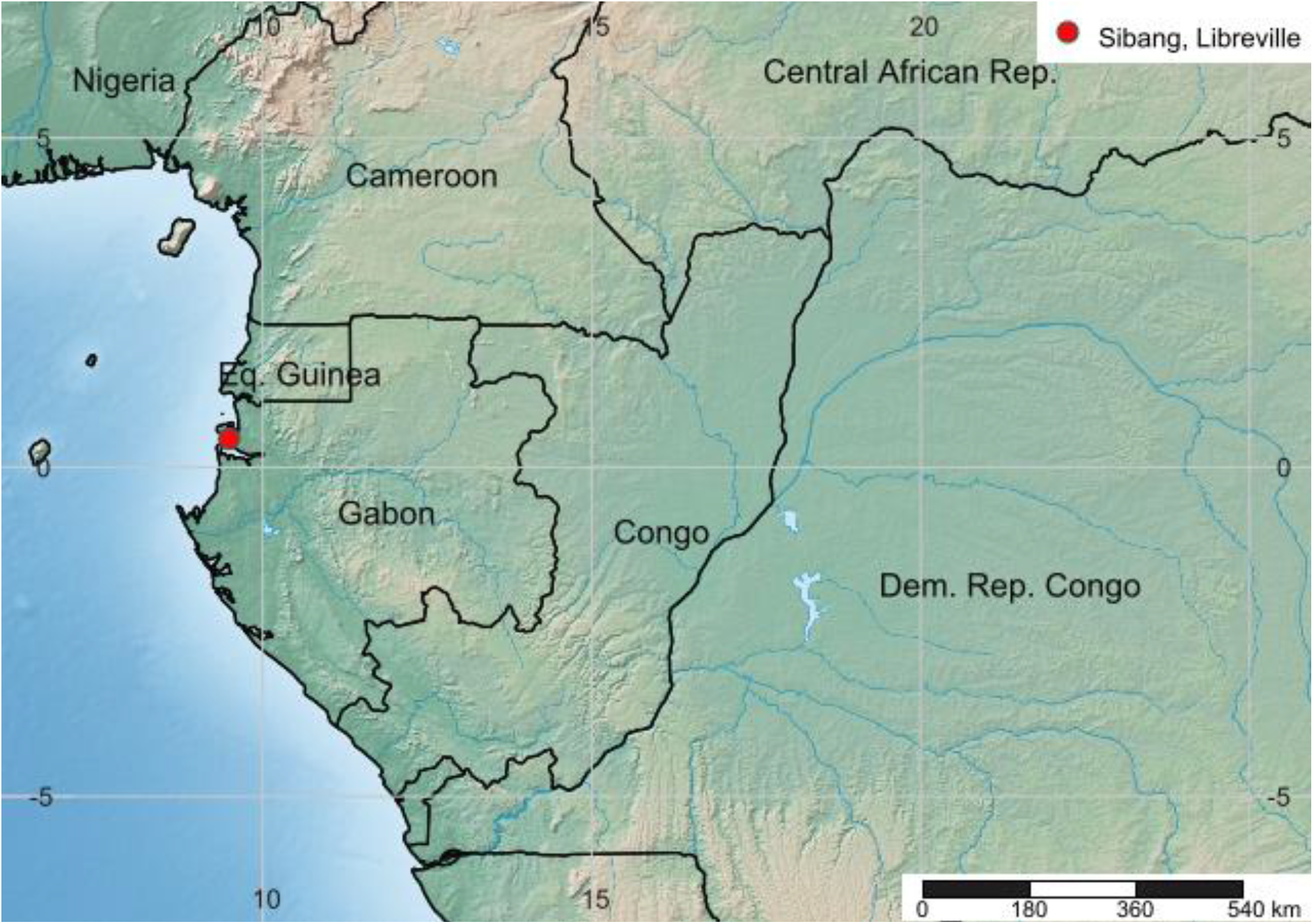
Global distribution of *Pseudohydrosme bogneri*. Red dot shows only known location, Libreville, Sibang.

## SPECIMENS EXAMINED

GABON, Libreville. Sibang Arboretum, fl. Oct. 1973, cultivated in Munich Botanical Garden, *Bogner* 662 (holotype K03822705; isotypes K03822706 spirit K37350.000, M six sheets, US03822705, US03822706); ibid, sterile, juvenile, 24 Oct. 1973, *Bogner* 640 (K000593312, spirit K29047.001).

## HABITAT

Known from one site in lowland rainforest on sandy loam, with *Aucoumea klaineana* Pierre (Burseraceae); c. 50 m alt.

## CONSERVATION STATUS

*Pseudohydrosme bogneri* is possibly extinct at its only known wild location. This is at Sibang, now in Libreville. Measured on Google Earth, the forest is approximately a square, c. 470 m N to S, and 420 m W to E, or about 0.25 km^2^ (grid reference: 0° 25’ 56.05”N, 9° 29’ 23.64” E, 49 m alt.). It is now completely surrounded by the dense urban settlement of Libreville which has expanded greatly in the last 60 years. In 1960, at independence, the population of Libreville was 32,000. Since then, it has expanded 200-fold to, in 2013, 703,904 (https://en.wikipedia.org/wiki/Libreville, accessed 19 Sept. 2020) and has a vastly greater footprint. Sibang Arboretum, the surviving patch of forest of a once much greater area, is now known as one of the top two tourist destinations in Libreville. This minute remnant of forest is probably the most visited by botanists in the whole of Gabon because it is immediately adjacent to the site of the current National Herbarium of Gabon, LBV (Cheek, pers. obs.). Because it has not been recorded here or anywhere else, in nearly 50 years since it was collected at Sibang by Bogner in 1973, it is possible that it no longer survives at this location. It was evidently rare at this location since Bogner reported seeing only a single plant (Bogner 1981). The Libreville area has been intensively studied botanically (Sosef *et al.* 2006, see discussion).

The area of occupancy is calculated as 4 km^2^ using the IUCN (2012) preferred gridcells of this size, and the extent of occurrence must therefore be of the same size following IUCN rules. Since less than 50 mature individuals are known in the wild, and because there is a single location, and since in any case the species is possibly extinct, we assess the species as CR D, B1+B2ab(iv), that is Critically Endangered (Possibly Extinct). In the unlikely but hoped for event that this conspicuous species should be found anywhere else in the vicinity of Libreville, it is likely to be threatened by human pressures since most of the population of Gabon is concentrated here, and abstraction of wood for fuel and construction from natural habitat is known to pose a threat (Lachenaud *et al*. 2013; Walters *et al*. 2016).

The peak decades of botanical collection in Gabon were at the end of the 20^th^ century. The Libreville region has the highest botanical specimen collection density in Gabon, with 5359 specimens recorded in digital format. It also has the highest level of diversity of both plant species overall and of endemics (Sosef *et al.* 2006). The coastal forests of the Libreville area are known to be especially rich in globally restricted species (Lachenaud *et al*. 2013; Gosline & Malecot 2011). These authors detail 19 species globally restricted to the Libreville area, of which eight have not been seen recently and which are possibly extinct (listed in Cheek *et al.* 2021). Among these is *Octoknema klaineana* Pierre, a rainforest tree “only collected in the immediate area of Libreville at the beginning of the 20^th^ century, and only once since.” Threats in the Libreville area have already resulted in the possible global extinction of these eight species, which rise to nine with the addition of *Pseudohydrosme buettneri* (Cheek *et al.* 2021) and ten when *P. bogneri* is included.

## PHENOLOGY

Flowering in late October, with leaf unfurling at anthesis, in Gabon; in cultivation in Europe, flowering in January. Fruiting unknown.

## ETYMOLOGY

The specific epithet honours Dr Josef Bogner, (1939 – 2020) respected and admired specialist in Araceae, who collected the only known material of this species in Oct. 1973.

## VERNACULAR NAMES & USES

Unknown.

## CULTIVATION

Josef Bogner appears to have taken two live collections of the new species in 1973 from Gabon to Munich Botanic Garden, both of which grew and flowered (since resulting cultivated material is preserved, see introduction above). In the case of *Bogner* 662 the plant flowered repeatedly (see above) and was propagated: at least one plant was given to the Tropical Nursery, Royal Botanic Gardens, Kew in 1976 where it was given the accession number 1976 – 789. The plant at Kew flowered (preserved specimens at Kew) but was recorded as dead in Jan. 1996 (Paul Rees in litt. to Cheek, March 2021). The plants at Munich also died (Andreas Groeger in litt. to Cheek, March 2021), at an unknown date but certainly after Jan. 1995 when one was photographed in flower (Fig. 1). The cause of the plant death, and the protocol for cultivation are thought to be unrecorded at both institutes. It is not known whether any plants survive in cultivation. We have not been able to trace any record of fruit or seed being produced. Seed production in *Pseudohydrosme* is not straightforward (Bogner & Hetterscheid 2013). Had seed been available, it is likely that this species would be widespread in cultivation since it is spectacular in flower (Fig. 1).

## NOTES

Searches for additional material of *Pseudohydrosme bogneri* have failed to find any specimens that definitely matches this taxon (but see note below). No additional specimens have been attributed to *Anchomanes nigritianus* in the literature apart from the type, *Bogner* 662 and *Bogner* 640 (the latter two specimens here described as *Pseudohydrosme bogneri*). In the checklist of the plants of Gabon only two taxa are recognised in *Anchomanes*, *A. difformis* and *A. giganteus* (Sosef *et al.* 2006). The last has a single specimen listed, *N. Hallé* 5481 (P). Given Bogner’s first identification of his specimens 662 and 640 as *Anchomanes giganteus* we examined the two sheets of this fruiting specimen carefully but found that they lack the diagnostic characters of *Pseudohydrosme bogneri*, rather they are *A. difformis*, since e.g. the spathe is c. 7cm wide. In *Pseudohydrosme bogneri* the spathe is 20 cm wide when spread out (see description above).

We examined other specimens filed as *A. difformis*, at P, WAG/L, M, MA and K in case material of the new species had been misidentified as this widespread and common taxon. However, no additional flowering specimens were found that matched the Bogner material. The two species are easily separable when in flower. *Anchomanes difformis*, apart from a much narrower spathe (see above) has the male zone of the spadix 2.5 – 4.5 times longer than the female zone (versus the male zone marginally shorter or equal to the female zone in the new species, *Pseudohydrosme bogneri)*. In addition, as in all *Anchomanes*, the spathe of *A. difformis* opens to the base, lacking a convolute tube which conceals the lower part of the spadix as is present in the new species, and is characteristic of *Pseudohydrosme*. However, a distinctive vegetative feature of *P.bogneri* are the long petiolules of the ultimate leaf segments, especially those most proximal to the petiole. In other species of the genus, and also of all *Anchomanes* known in West-Central Africa, the ultimate leaf segments are either sessile, or with a very short petiolule <0.1 – 0.2 cm long. One sterile specimen *N.Hallé* 2448 (P), previously identified as *Anchomanes difformis*, exhibited petiolules that were 0.5 – 0.15 cm long. Since not so long as the petiolules of *P. bogneri* (0.5 – 2.5 cm long), we hesitate to identify it as such without an inflorescence. Collected in Gabon at “Abanga” in Moyen-Ogooué Province, we advise that this area be searched in late October for flowering material to determine the identity of the taxon concerned. It might prove to be a further new taxon, or conceivably extend the morphological and geographical range of *Pseudohydrosme bogneri*.

### Bogner’s placement in *Anchomanes*

Bogner’s placement of his collections 640 and 662 in *Anchomanes* is easy to understand for two reasons: 1) the long spiny, terete peduncle, far exceeding the spathe in length, has hitherto been seen only in this genus, at least in Guineo-Congolian Africa; 2) as reported in the introduction, the new species bears its inflorescence at the same time that the leaf is unfurling, as usual in *Anchomanes*. In other *Pseudohydrosme* species, the leaf does not usually begin to emerge from the ground until some weeks after flowering is finished. There is a natural human tendency on encountering an unfamiliar species to place it with a taxon that is already recognised.

## DISCUSSION

The discovery of a fourth, albeit anomalous, species of *Pseudohydrosme* in Libreville is congruent with existing knowledge of the genus (Cheek *et al.* 2021). Of the three species previously known, two were discovered in what is now Libreville, and one of these, *P. buettneri* Engl. which is apparently now extinct, was a point endemic. Most of the records of the other surviving species, *Pseudohydrosme gabunensis*, were derived from either Libreville or the immediate area. The third species *Pseudohydrosme ebo* Cheek, is a geographical outlier, found far to the north in the Cross-Sanaga Interval (Cheek *et al.* 2001), a separate phytogeographic area c. 500 km to the North. With the discovery of this fourth species of *Pseudohydrosme*, Libreville is confirmed as the centre of diversity for *Pseudohydrosme*.

The morphology of the new species is so discordant from the other species of the genus, due to the long peduncle, and the long spadix in proportion to the spathe, that it challenges the existing generic delimitation. Yet this is consistent with the high morphological divergence between the two first described species, which is so great that they were previously placed in separate genera (Brown in Thistleton Dyer 1901). The large morphological differences between the species in the Libreville area suggest that not only did the genus originate there but that it is not actively evolving since the morphological gaps between the taxa are so large. Rather, in complete contrast to *Anchomanes, Pseudohydrosme* seems to be a relictual genus, supported by the fact that all its species are rare, localised, and highly threatened with extinction, or likely already extinct (*Pseudohydrosme buettneri*). The least threatened species of the genus is *Pseudohydrosme gabunensis* which was assessed as Endangered, EN B2ab(ii,iii) (Lovell & Cheek 2020). Even the geographically disjunct Cameroonian *Pseudohydrosme ebo*, the exception since it is morphologically close to *P. gabunensis*, appears restricted to a very small area and is Critically Endangered (Cheek *et al.* 2021).

## CONCLUSIONS

Such cases as *Pseudohydrosme bogneri* underline the urgency for identifying and publishing species while they still survive. Threats to such new discoveries for science are clear and current, putting these species at high risk of extinction, if they are not extinct already as may be the case here. About 2000 new species of flowering plant have been discovered by science each year for the last decade or more (Cheek *et al.* 2020), adding to the estimated 369 000 already known (Nic Lughadha *et al.* 2016), although the number of flowering plant species known to science is disputed (Nic Lughadha *et al.* 2017). Only 7.2% have been assessed and included on the Red List using the IUCN (2012) standard (Bachman *et al.* 2019), but this number rises to 21–26% when additional evidence-based assessments are considered, and 30–44% of these assess the species as threatened (Bachman *et al.* 2018). Newly discovered species, such as that reported in this paper, are likely to be threatened, since widespread species tend to have been already discovered. There are notable exceptions to this rule (e.g. *Vepris occidentalis* Cheek (Cheek *et al.* 2019) a species widespread in West Africa from Guinea to Ghana). Generally, it is the more localised, rarer species that remain undiscovered. This makes it all the more urgent to find, document and protect such species before they become extinct. Until species are delimited, described and known to science, it is difficult to assess them for their IUCN conservation status and so the possibility of protecting them is reduced (Cheek *et al.* 2020). *Pseudohydrosme bogneri* seems to be the tenth extinct plant species from the Libreville area (see discussion above), which is likely to be the area with the greatest number of plant species extinctions in Gabon. Documented extinctions of plant species are increasing, e.g., in neighbouring Cameroon, *Oxygyne triandra* Schltr. and *Afrothismia pachyantha* Schltr. have recently been shown to be globally extinct (Cheek *et al.* 2018a; Cheek & Williams 1999, Cheek *et al.* 2019), and as seems to be the case with *Pseudohydrosme bogneri*, there are also examples of species that appear to have become extinct even before they are known to science, such as *Vepris bali* Cheek (Cheek *et al.* 2018b). In all cases anthropogenic pressures have been the cause of these extinctions. There is no Red Data Book for the plants of Gabon, but in that for neighbouring Cameroon it was found that of the 815 threatened species recognised, most were threatened by habitat clearance and/or degradation, following logging, especially of forest for small-holder and plantation agriculture e.g. oil palm (Onana & Cheek 2011). In Gabon, on the whole, natural habitat is much more intact, and much better covered by a National Park network than in Cameroon. However, delimitation in Gabon of the highest priority areas for plant conservation as Tropical Important Plant Areas (TIPAs, using the revised IPA criteria set out in Darbyshire *et al.* (2017)) as is in progress in Cameroon, should help to avoid future global extinctions of range-restricted endemic species such as *Pseudohydrosme bogneri*.

## Acknowledgements

We thank Marcelo Sellaro of the Tropical Nursery, RBG, Kew for introducing us to Dr. Andreas Groeger and Dr. Günter Gerlach of Munich Botanic Garden. We thank the latter for providing and permitting our use of his excellent photos of the new species in this paper. Dr Andreas Fleischmann of M kindly provided images of the herbarium specimens requested. Andrea Hart of the Natural History Museum, London offered help tracking original material of *Anchomanes nigritianus*. Paul Rees tracked down records at the Tropical Nursery, Kew Janis Shillito typed much of the manuscript. Alastair Hay is thanked for inspiring this project. We owe Simon Mayo thanks for urging us to investigate the spirit collection at K for *Anchomanes*.

